# Loss of cell cycle control renders cells nonresponsive to local extrinsic differentiation cues

**DOI:** 10.1101/720276

**Authors:** Kara L. Cerveny, Ingrid Tower, Dayna B. Lamb, Avery Van Duzer, Hannah Bronstein, Olivia Hagen, Máté Varga

## Abstract

**Objective and approaches:** Aberrantly proliferating cells are linked to a number of diseases including cancers and developmental defects.To determine the extent to which local extrinsic signals contribute to or ameliorate mutant cell behaviors, we examined survival and differentiation of mutant cells in wild-type retinal environments by generating chimeric zebrafish embryos comprised of unlabeled host cells and GFP-labeled neural progenitor donor cells. In addition, we examined the fate of retinal progenitor cells when *cdkn1c*, a cyclin dependent kinase inhibitor, was induced in clones within wild-type and *hdac1* mutant retinae.

**Results:** We found that seven of the ten mutants examined exhibited apoptosis when grafted into wild-type tissue, with cells from two slowly cycling mutants, *elys* and *emi1*, noticeably differentiating in a wild-type environment. Observations of the one hyperproliferative mutant, *hdac1*, revealed that these mutant cells did not appear to die or differentiate but instead survived and formed tumor-like rosettes in a wild-type environment. Ectopic expression of *cdkn1c* was unable to force cell cycle exit and differentiation of the majority of *hdac1* mutant cells.

**Conclusions:** Together, these results suggest that although a wild-type environment rarely encourages cell cycle exit and differentiation of neural progenitors with cell cycle defects, wild-type survival signals may enable hyperproliferative progenitor cells to persist instead of die.

## Introduction

Strict control of proliferation, cell cycle exit, and differentiation underlies the formation, maintenance, and repair of nervous tissues of appropriate size and composition (Morales and Mira, 2019; Schmidt et al., 2013; Urbán and Guillemot, 2014). Mutations in genes that control these processes are linked to cancer and overgrowth syndromes (Fruman et al., 2017; Santamaria and Ortega, 2006). Proliferation is intrinsically-controlled through the precisely timed synthesis and degradation of cyclins, proteins that activate cyclin-dependent kinases (CDKs) to propel the cell through sequential phases of DNA replication, growth, and mitosis. Cyclin-dependent kinase inhibitors (CKIs), cell cycle phase specific ubiquitin ligases, and other regulatory proteins also govern cell cycle progression, ensuring sufficient growth, accurate DNA replication, and equal chromosome segregation during each phase of the cell cycle (Murray, 2004; Santamaria and Ortega, 2006). Cells employ a variety of intrinsic mechanisms to ensure that the decision to proceed from one phase of the cell cycle to the next is appropriate. For example, cells transitioning from DNA replication to chromosome segregation survey their DNA for damage, either satisfying a molecular checkpoint and passing into the next phase or activating checkpoint controls that initiate DNA repair or, if the damage is extensive, cell death (Murray and Carr, 2018).

In addition to the well-studied checkpoint control that is shared by all cells, neural progenitors and neurons themselves require additional cell cycle control mechanisms to ensure appropriate proportions of distinct types of neurons and glia (Frade and Ovejero-Benito, 2015). For example, differentiated neurons in the vertebrate central nervous system have been found to contain chromosomal abnormalities including tetraploidy consistent with proliferation defects, suggesting that differentiation rather than death of defective cells is possible in particular tissues (e.g., (Zupanc et al., 2009); reviewed in (Frade and Ovejero-Benito, 2015)). Furthermore, several studies of zebrafish eyes have shown that a differentiated retinal environment can support survival and differentiation of aberrantly proliferating cells that typically die (Cerveny et al., 2010; Link et al., 2000). In some contexts, differentiation is not incompatible with continued proliferation as horizontal cells in the retina have been shown to proliferate and contribute additional neurons to the developing retinal circuitry (Godinho et al., 2007). These observations raise the possibility of central nervous system (CNS)-specific environmental control over proliferation and differentiation decisions.

Extrinsic input into proliferation and differentiation decisions, especially in multicellular organisms, is not unexpected. In the vertebrate retina, Müller glia can be stimulated to re-enter proliferation and support retinal growth and regeneration in response to a number of extrinsic factors including insulin growth factors, FGF, and TNFα (e.g.,(Conner et al., 2014; Wan et al., 2014). In addition to secreted proteins, exogenous teratogens such as ethanol have been shown to perturb proliferation and differentiation, especially in the developing nervous system, by impinging on extrinsically-regulated signaling pathways (e.g., Kashyap et al., 2007; Muralidharan et al., 2018). Extrinsically-regulated signaling pathways are also known to stimulate cell cycle exit and differentiation, with pathways like Notch-Delta balancing proliferation and differentiation decisions of neighboring cells through lateral-inhibition type mechanisms (Louvi and Artavanis-Tsakonas, 2006). Recent analysis of medaka retinae provide evidence that Notch pathway activation appears to be limited to a subset of progenitor cells and is required for generating eyes with appropriate proportions of neurons and glia (Pérez Saturnino et al., 2018). In addition, locally secreted molecules can encourage cell cycle exit and differentiation of cycling progenitors during development. For example, the spatial and temporal pattern of Hedgehog pathway activity, in both invertebrate and vertebrate eyes, ensures timely cell cycle exit and differentiation of retinal progenitors (García-Morales et al., 2019; Locker et al., 2006; Masai et al., 2005; Neumann and Nuesslein-Volhard, 2000). In addition, retinoic acid (RA) can promote cell cycle exit and neuronal differentiation, biasing cells toward various neuronal fates both in vivo (eg., Hyatt et al., 1996; Stevens et al., 2011; Valdivia et al., 2016) and in tissue culture (e.g.,Estephane and Anctil, 2010; Kelley et al., 1999). Whether these extrinsic signals could be used to force cells to differentiate rather than die or undergo hyperproliferation in disease contexts is an open question.

To systematically examine whether wild-type extrinsic signals could promote survival and differentiation of aberrantly cycling cells, this study first identified and assessed an arbitrary sample of 10 mutant zebrafish lines (see Table 1). It then generated chimeric retinae to assess behaviors of those mutant cells in a wild-type context. Finally, it investigated whether ectopic expression of an intrinsic cyclin dependent kinase inhibitor, Cdkn1c, which is linked to cell cycle exit and differentiation, could force differentiation of hyperproliferative cells in developing retinae. The data presented here confirm that neurotypical cell proliferation and differentiation often require both intrinsic factors and extrinsic cues. In addition, the data indicate that cells carrying mutations in genes encoding the cell cycle machinery do not appear to be susceptible to differentiation cues from the local environment.

**Table 1.** Cell cycle mutants examined by chimeric analysis in the zebrafish retina

| **mutant** | **phenotype linked to cell cycle defect** | **phenotypes of mutant cells transplanted into WT retinæ** | **Reference** |
| --- | --- | --- | --- |
| *cdk1<sup>hi3235Tg</sup>* | stall in G1, G1/S, S phases, apoptosis | apoptosis | (Amsterdam et al., 2004); this study |
| *dtl<sup>hi3627Tg</sup>* | arrest in late S/early G2, apoptosis | apoptosis | (Sansam et al., 2010); this study |
| *ele<sup>ty77e</sup> (slbp1)* | stall in G1/S, apoptosis | apoptosis | (Turner et al., 2019) |
| *emi1<sup>hi2648Tg</sup> (fbxo5)* | primarily arrest in G2/M, apoptosis. | some differentiation but also apoptosis | (Rhodes et al., 2009; Riley et al., 2010; Zhang et al., 2008); this study |
| *flo<sup>ti262c</sup> (ELYS)* | cycle slowly, stalling in either G1/S or G2/M | survival and differentiation | (Cervený et al., 2010; Davuluri et al., 2008); this study |
| *gins2* | Delayed/prolonged S phase, apoptosis | apoptosis | (Varga et al., in preparation.) |
| *hdac1<sup>hi1618Tg</sup>* | unable to exit the cell cycle; hyperproliferate and do not differentiate | survival and proliferation | (Yamaguchi et al., 2005; Zhou et al., 2011); this study |
| *mcm5<sup>m850</sup>* | prolonged S phase, apoptosis | apoptosis | (Ryu et al., 2005); this study |
| *cmp<sup>s819</sup> (ssrp1a)* | arrest in S phase, apoptosis. | quiescence and apoptosis | (Koltowska et al., 2013); this study |
| *rbbp6<sup>hi2993Tg</sup>* | predicted to arrest in G1/S, apoptosis | apoptosis | (Amsterdam et al., 2004); this study |

## Results and Discussion

To test whether a wild-type environment could encourage the differentiation of neuronal progenitor cells with impaired cell cycle control, we created chimeric zebrafish embryos containing mutant cells in wild-type retinae. We examined all retinae at 3 days post-fertilization (dpf) and found that out of the ten strains that we tested, nearly all cell cycle deficient cells failed to alter their behavior in a wild-type environment (Table 1). Consistent with previous reports, retinal progenitor cells (RPCs) in *cdk1, dtl, ele, emi1, elys, gins2, mcm5, ssrp1a*, and *rbbp6* deficient cells appeared to die by apoptosis (Table 1; references therein) as we observed pyknotic nuclei throughout the developing retinal neuroepithelium at 48 and 72 hours post fertilization (hpf) of all embryos (data not shown). When mutant cells were transplanted into wild-type hosts, the only clear evidence of mutant cell differentiation was found in chimeric retinae containing *emi1* or *elys* homozygous mutant cells (Figure 1C-H). All of the other mutant cells exhibit blebbing and fragmentation when integrated into wild-type retinae, consistent with cell-autonomous apoptosis (e.g., Figure 1I-J; Table 1).

**Figure 1.**
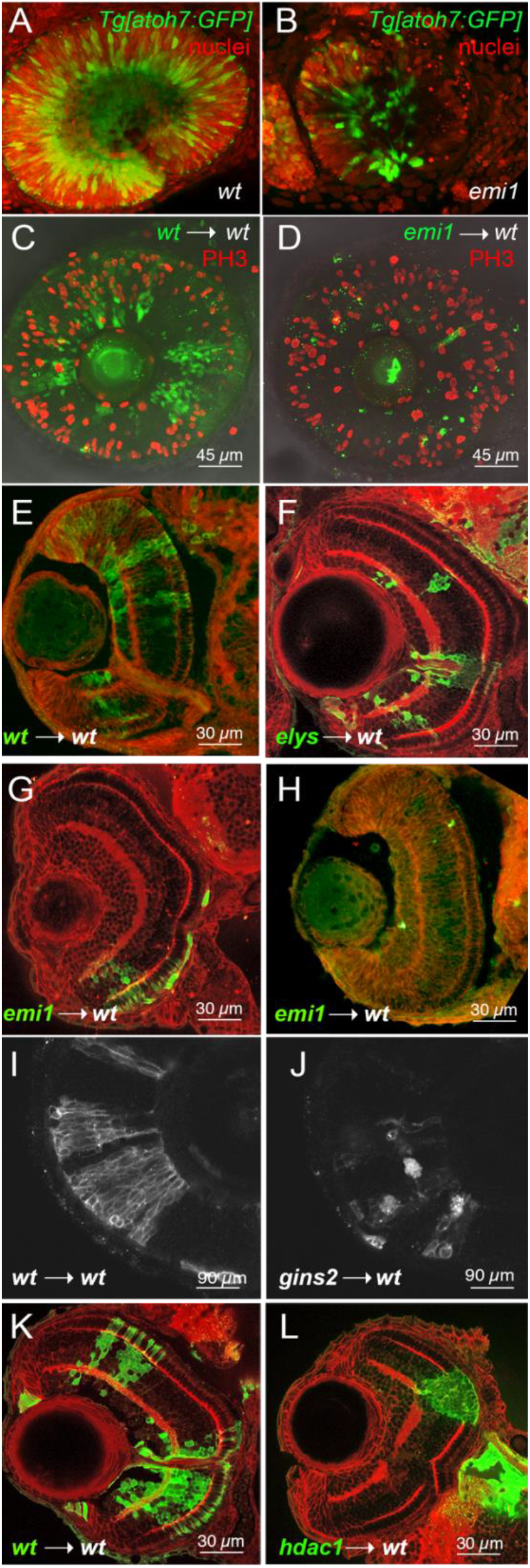
Comparison of differentiation behavior of wild-type, *emi1^−/−^*, and *elys^−/−^* neuronal progenitors in developing zebrafish retinae. A-B Lateral maximum intensity projection of 50 hpf retinae from *Tg[atoh7:GFP]* embryos showing progression of neurogenic gene expression (green) stained with sytox orange (red) to highlight nuclei. C-D Lateral views of maximum intensity projection of 3 dpf chimeric wild-type retinae containing GFP-labelled wild-type (C) or *emi1* mutant (D) cells and immunostained for phosphohistone H3 (PH3, red). E-H Representative images of frontal cryosections of 3 dpf chimeric wild-type retinae containing GFP-labelled wild-type (E), *elys* mutant (F), or *emi1* mutant (G, H) cells immunostained for GFP (green cells from donor embryo) and B-catenin (red cell boundaries and plexiform layers). I-J Lateral views of single z-slice of ventral-nasal region of 3 dpf chimeric wild-type retinae containing wild-type (I) or *gins2* morphant (H) cells labeled with membrane-targeted RFP. K-L Representative images of frontal cryosections of 3 dpf chimeric wild-type retinae containing GFP-labelled wild-type (K) or *hdac1* mutant (L) cells immunostained for GFP (green cells from donor) and B-catenin (red) to mark cell boundaries and plexiform layers.

Previous reports have shown that zebrafish embryos carrying homozygous mutations in *emi1* (also known as *fbxo5)* and *elys* still exhibit some neuronal differentiation (Cerveny et al., 2010; Riley et al., 2010; Zhang et al., 2008), potentially due to maternal inheritance of these mRNAs or stability of the protein. For example, a small but significant fraction of *emi1* mutant RPCs still express the neurogenic gene *atoh7*, exit the cell cycle, and differentiate into retinal ganglion cells in *emi1* mutant eyes (Figure 1A-B). The same has been shown for *elys* mutants (Cerveny et al., 2010; Davuluri et al., 2008). Interestingly, we found that all *elys* mutant cells transplanted into wild-type eyes appeared to differentiate (Figure 1F, n = 19 transplants;(Cerveny et al., 2010)) whereas *emi* mutant cells transplanted into wild-type eyes either survived and differentiated (Figure 1G, n = 10/34 transplants) or appeared to quiesce or be lost due to cell death (Figure 1H, n = 24/34 transplants). Because we controlled for the number of cells transplanted into each embryo, survival and differentiation of *emi1* cells in a wild-type environment were possibly influenced by the location of the transplanted cells, slight differences in age of the host embryo at time of analysis, or stochastic fluctuations in gene expression in the transplanted cells or host embryos.

The difference in susceptibility of *emi1* and *elys* mutant cells to differentiation factors from the wild-type environment may be explained by the distinct functions of these mutated genes. For example, the *emi1* gene encodes a protein that directly participates in the cell cycle by acting as both a substrate for and inhibitor of the anaphase promoting complex (APC/C) (Cappell et al., 2018), whereas the *elys* gene encodes a large scaffold protein required for nuclear pore formation and possibly chromatin organization (Rasala et al., 2006). It is therefore tempting to speculate that differences in epigenetic regulation may underlie the complete differentiation of *elys* cells transplanted into a wild-type retina. It is also important to note that of all the mutants we tested, only *elys* does not carry a mutation in a gene directly linked to cell cycle progression or regulation.

Our transplant studies also confirmed previous reports that mutations in histone deacetylase 1, *hdac1*, are linked to cell autonomous hyperproliferation in the retina ((Stadler et al., 2005; Yamaguchi et al., 2005), Figure 1K-L). When we examined *hdac1* mutant cells integrated into wild-type chimeric retinae at 3 dpf, a point at which apoptotic cells are found scattered throughout the *hdac1* mutant retinae (Supplemental Figure 1), we did not observe pyknotic nuclei or cell blebbing, two other key hallmarks of apoptosis. This finding raises the possibility that a wild-type retinal environment supports the survival of these hyperproliferative cells but it does not alter their uncontrolled proliferation.

Intrigued by the survival and continued proliferation of *hdac1* mutant cells in wild-type embryos, we further analyzed wild-type retinae containing *hdac1* mutant clones at 4 dpf. Wild-type cells transplanted into a wild-type environment appeared to differentiate as expected and wild-type cells in an *hdac1* environment also exhibited hallmarks of differentiation, forming clones that contained cells with typical photoreceptor and interneuron morphologies (Figure 2A,C). Small clones of wild-type cells did not appear to force neighboring mutant cells to differentiate (Figure 2C). Interestingly, we found that *hdac1* mutant cells formed rosettes in wild-type retinae, reminiscent of early tumor formation (Figure 2B). Not only do these rosettes persist in wild-type eyes, but they also appear to disrupt adjacent retinal architecture, breaching the outer boundary of the apical surface (Figure 2B’’) and interrupting the inner plexiform layer. Moreover, these clones of *hdac1* mutant cells appear to disrupt the lamination and possibly differentiation of neighboring wild-type cells (Figure 2B’’). In 9 of 14 chimeric wild-type retinae containing *hdac1* mutant cells, we observed similar phenotypes of disrupted lamination and apical boundaries as shown in Figure 2B’’, raising the possibility that persistently cycling cells can alter the organization of wild-type tissues. We believe that these studies establish the chimeric retina approach as a potentially powerful way to study and understand the effects of nascent tumors on surrounding neuroepithelial tissues.

**Figure 2.**
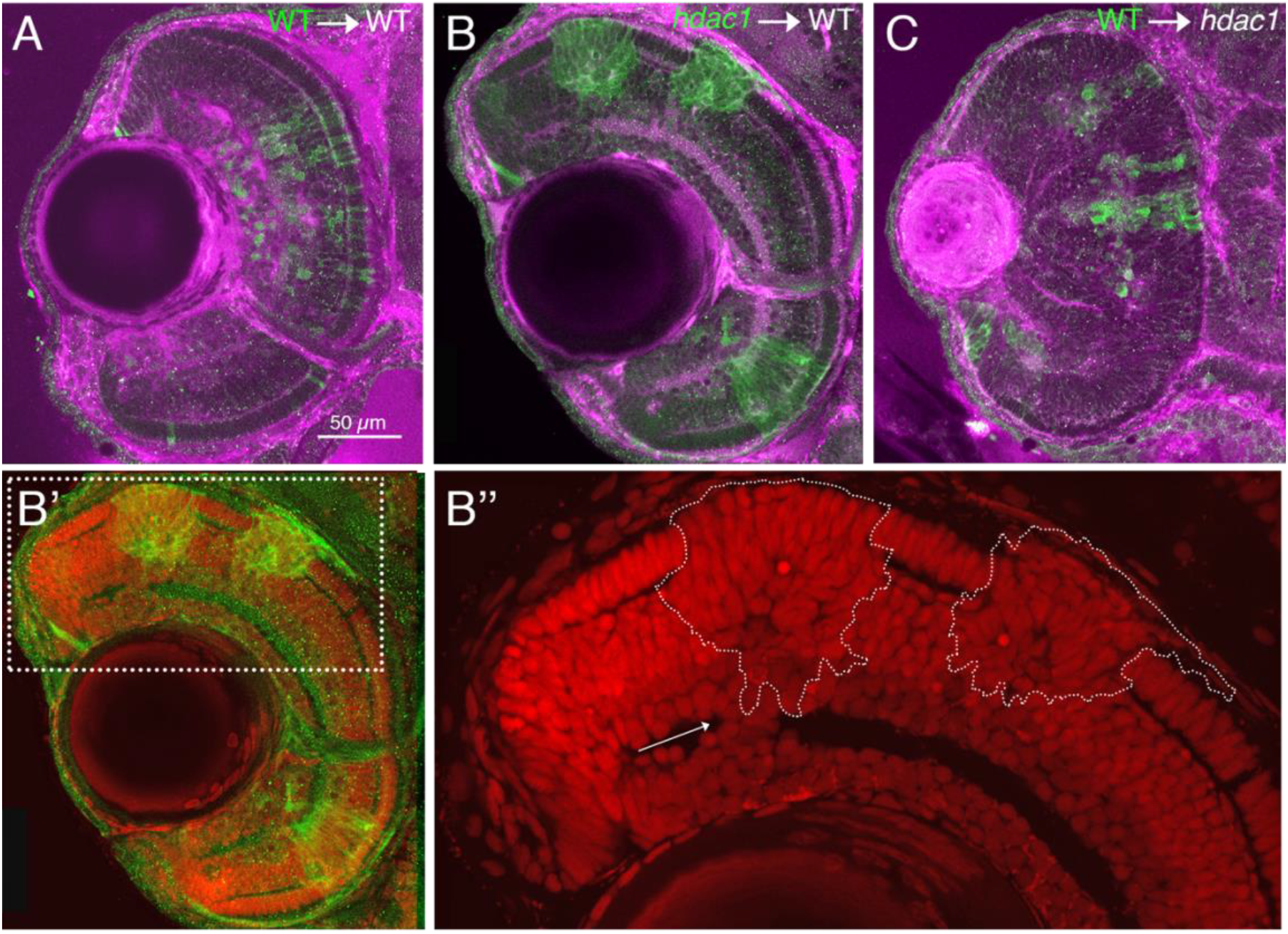
*hdac1* mutant cells form rosette-like structures that invade surrounding tissues when transplanted into WT retinae and alter lamination and organization of adjacent wild-type cells. A-C Frontal cross-sections of 4 dpf chimeric retinae when GFP-expressing wild-type (A, C) and *hdac1* mutant (B) cells were transplanted into wild-type (A, B) or *hdac1* mutant (C) host eyes immunostained for ZO-1 (purple) and GFP (green). B’-B’’ Same cross section as in B, this time showing transplanted cells (green) and DAPI-stained nuclei (red). Boxed region ~2X zoom shown in B’’. Arrow indicates wild-type cells disrupting the inner plexiform layer within this chimeric retina and *hdac1^−/−^* clone outlined with dashed line.

A previous study showed that *hdac1* mutant retinae are likely hyperproliferative because they fail to express key cell cycle exit genes including cyclin dependent kinase inhibitors (Yamaguchi et al., 2005). We asked whether *hdac1* mutant cells could be forced to exit the cell cycle and differentiate by inducing expression of *cdkn1c* (also known as *p57*) in *hdac1* mutant retinae. Contrary to previous reports showing that over-expression of a different cyclin dependent kinase inhibitor, *cdkn1b* (also known as *p27*), could induce cell cycle exit and differentiation of *hdac1^−/−^* retinal progenitor cells (e.g., (Ohnuma et al., 1999; Yamaguchi et al., 2005), we found that mosaic expression of *cdkn1c* from a heat-shock-inducible promoter from 26-28 hpf was rarely associated with differentiation of *hdac1* mutant RPCs (Figure 3C-E). Specifically, clones expressing *cdkn1c* always revealed morphologies typical of retinal neurons, most obviously photoreceptors, bipolar cells, and ganglion cells in wild-type eyes (e.g., Figure 3C; Supplemental Figure 2C) whereas nearly all *cdkn1c*-positive cells in *hdac1* mutant eyes appeared neuroepithelial (e.g., Figure 3D; Supplemental Figure 2B). In contrast to our observations of *hdac1^−/−^* wholemount eyes, our observations of thin cryosections of eyes with clones of *cdkn1-*expressing cells revealed a small minority of *hdac1^−/−^* clones (2 clones out of 26 clones in 9 eyes) that contained some cells with neuronal morphologies (Supplemental Figure 2A). These same eyes, as well as most other eyes, contained clones that did not exhibit neuronal morphology. Based on the GFP intensity in the *hdac1^−/−^* cells that appeared to differentiate, it is possible that either extremely high levels of Cdkn1c and/or slight differences in developmental timing of *cdkn1c* expression may support retinal progenitor cell differentiation in the *hdac1^−/−^* background.

**Figure 3.**
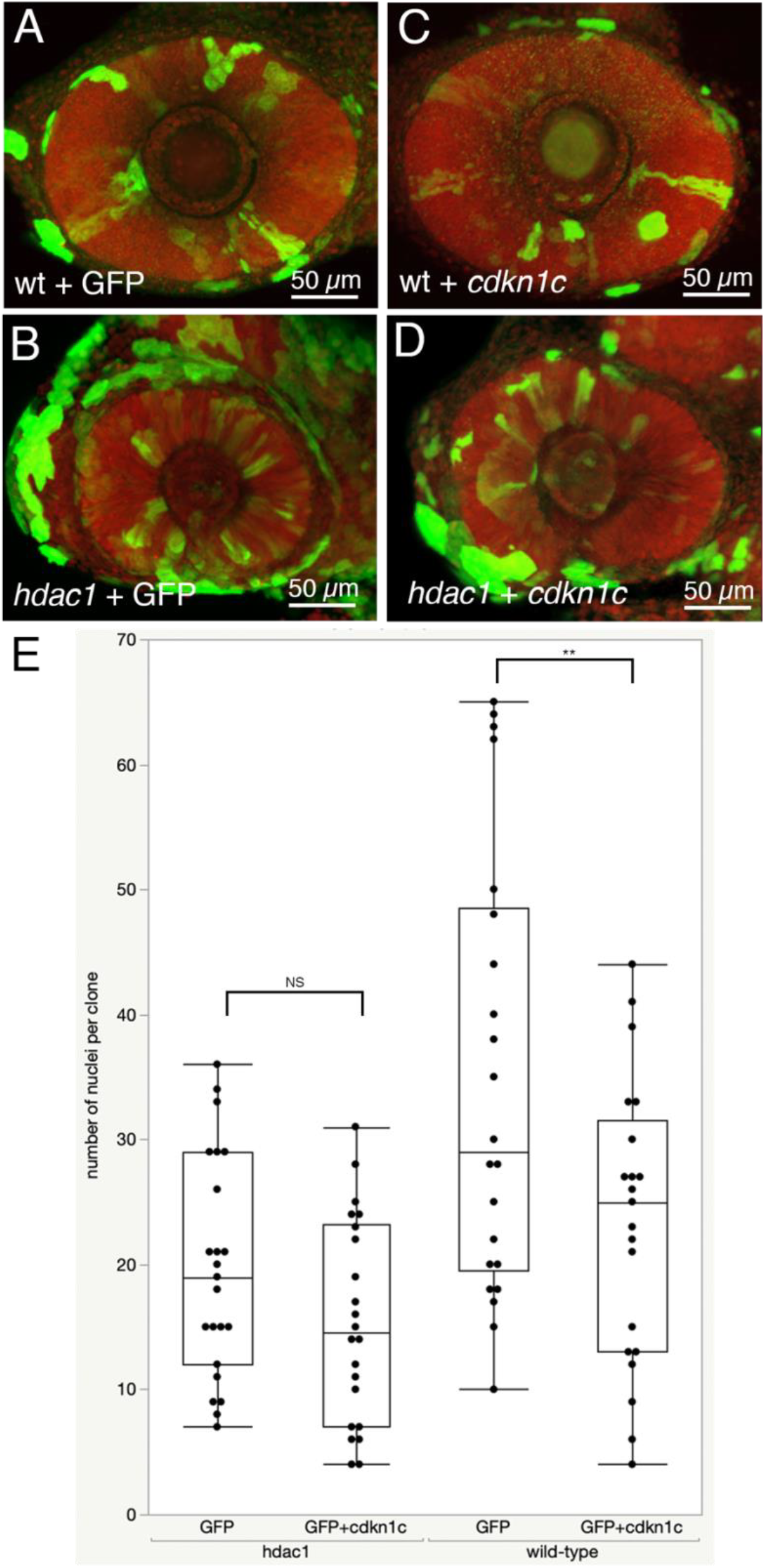
Forced expression of *cdkn1c* does not significantly alter proliferative behavior of *hdac1* mutant cells. A-D Lateral views of maximum intensity z-projections showing nuclei (red) and heat-shock induced GFP (A-B, green) or *cdkn1c*-postive clones (C-D, green) in wholemount 60 hpf embryos with genotype indicated. E Box plot overlaid on all data points for number of nuclei per clone in GFP and *cdkn1c* containing clones in either *hdac1* and wild-type sibling retinae. NS, not significant with p-value of p=0.1008; **, significant with p-value of 0.0174; Welch’s t-test.

To further explore how Cdkn1c activation altered *hdac1* and wild-type cells, we used precisely timed heat-shock to induce expression of either GFP alone or Cdkn1c and GFP in 28 hpf embryos and then measured the size of clones by counting the number of nuclei in GFP-labelled retinal clones in 60 hpf embryos. Based on clone size, wild-type retinal progenitor cells appeared to be more likely to exit the cell cycle and differentiate when *cdkn1c* was induced, but *hdac1* clones with and without *cdkn1c* appeared similar. Specifically, we found that control (GFP only) clones in wild-type retinae contained an average of 34.5 nuclei whereas *cdkn1c*-expressing clones in wild-type retinae contained an average of only 23.3 nuclei per clone (Figure 2E; n=22 clones, 8 eyes for each wild-type sample; p=0.0174, Welch’s t-test, 95% CI). These data suggest that high levels of *cdkn1c* can encourage cell cycle exit and differentiation of proliferating retinal progenitor cells. In contrast, control clones in *hdac1* mutant retinae contained an average of 19.6 nuclei and were not much larger than *cdkn1c-*expressing clones in *hdac1* mutant retinae, which had an average of 15.4 nuclei (Figure 2E; n=23 clones, 9 eyes for each *hdac1^−/−^* sample; p=0.1008, Welch’s t-test, 95% CI). Interestingly, clone size in *hdac1* embryos did not differ from *cdkn1c*-positive clones in wild-type retinae, suggesting that *hdac1*-linked hyperproliferation is not linked to an increase in cell cycle speed but a failure to ever exit the cell cycle. Taken together with our observations of limited differentiation in *hdac1* mutant cells containing induced *cdkn1c*, these data support the idea that Hdac1 activity likely contributes to expression of a number of gene products that ultimately work together to efficiently promote cell cycle exit and differentiation.

## Conclusions and Limitations

Based on our analyses of cell cycle mutant cells in wild-type retinal environments, we suggest that cell cycle machinery and/or cell cycle control components are intrinsically required for cells to respond to local extrinsic differentiation cues. We observed that nearly all cells carrying mutations in genes directly controlling the cell cycle were not encouraged to differentiate by a wild-type environment. The only mutant with direct links to cell cycle control that appeared to have both autonomous and non-autonomous behaviors in a wild-type environment was *emi1*. A recent study shows that cell cycle progression, and by extension, cell cycle exit, can be differentially influenced by levels of Emi1 protein, with low levels of Emi1 linked to quiescence and high levels linked to S-phase entry and robust DNA replication (Cappell et al., 2018). We speculate that small fluctuations in *emi1* mRNA is one possible explanation for the variable behavior we observe when *emi1* mutant cells are transplanted into wild-type retinae.

In this study and in previous studies (Cerveny et al., 2010), we observed that cells carrying mutations that likely have secondary effects on cell cycle progression (e.g., *elys*) can reliably survive and differentiate in a wild-type environment. The gene that is mutated in *elys* encodes a key component of the nuclear pore complex that possibly links nuclear pore assembly, nucleus size and DNA replication with chromatin organization (Gillespie et al., 2007; Jevtić et al., 2019; Zierhut et al., 2014). Similar to *elys* mutants, cells carrying a mutation in the chromatin-remodeling gene *brg1* can be encouraged to exit the cell cycle and differentiate in a wild-type environment (Gregg et al., 2003; Link et al., 2000). We therefore propose that global changes in genome organization underlie the phenomenon of environmentally enforced differentiation.

Not all types of genome organization defects are equivalent, however, as *hdac1* mutants appeared to act in a cell-autonomous manner, continuing to proliferate in a wild-type environment. We were, however, unable to follow the long-term fate of the *hdac1* mutant transplants past 4 dpf and so were unable to establish whether these clusters of cells evolve into full-blown retinal tumors. It would also have been interesting to further explore how the surrounding wild-type tissues and cells alter their organization and behavior.

Our observations and those previously examining the effect of *hdac1* mutation in the CNS (Schultz et al., 2018; Stadler et al., 2005; Yamaguchi et al., 2005) run counter to the vast collection of data showing that pharmacological inhibition of histone deactylase (HDAC) activity prevents over-proliferation, especially of cancerous cells (e.g., Li and Seto, 2016). Both laboratory and clinical studies provide evidence that inhibition of HDAC activity blocks proliferation and often promotes apoptosis of proliferating, oncogenic cells. In differentiated brain tissue, however, inhibition of HDAC activity is linked to neuronal survival, especially in neurodegenerative disease models (e.g., Didonna and Opal, 2015). Histone deacetylases not only remove acetyl groups from histones but are also known to deacetylate other targets including the tumor suppressor p53. Interestingly, one study provides evidence that maintenance of acetylation at specific lysine residues in p53 prevents its association with chromatin specifically in neurons (Brochier et al., 2013). Whether this type of regulation for p53 occurs only in differentiated neurons or also in neuronal progenitors, such as the RPCs examined in this study, is unknown.

## Methods

### Zebrafish lines

Eggs were collected by natural spawning, raised at either 25°C or 28.5°C in E3 embryo medium (Nüsslein-Volhard, C. and Dahm, R., 2002) and staged according to Kimmel et al., 1995. After gastrulation and before 24 hours post fertilization, embryos were cultured in 0.003% phenylthiourea (PTU, Sigma) in E3 to prevent pigment formation. Lines used in this study and references are listed in Table 1. Adult zebrafish were cared for with protocols approved by the Reed College IACUC.

### Cell transplants

Similar to previously published studies (e.g., (Cerveny et al., 2010; Turner et al., 2019), donor embryos were injected at the 1-cell stage with ~20 ng of GFP mRNA synthesized from linearized pCS2-GFP or membrane-targeted RFP mRNA synthesized from linearized pCS2-mCherry with the T7 mMessage mMachine kit (Ambion) according to manufacturer’s instructions. Host and donor embryos were grown at 28.5°C until sphere stage (approximately 4 hours post-fertilization) and then 10-15 fluorescently labelled cells were removed from donor embryos and transplanted into the animal pole of unlabled host embryos. Donor and host embryos were incubated overnight at 28.5°C. All embryos were screened and E3 was exchanged for PTU in E3. Donors were identified by visual inspection and by PCR and restriction digest mediated genotyping. Genotyping protocols for each line can be found at Zebrafish International Resource Center (ZIRC.org) and in relevant references (see Table 1). For *gins2* experiments, 1-cell stage embryos were first injected with ~1 nl of 1 mM gins2 morpholino (Gene Tools, Philomath, OR; 5’-GGGGTGAGTCAATTTATAATCTAC-3’), a dose that phenocopies *gins2*^−/−^ mutants (Varga et al., in preparation) and then injected with ~10 ng of membrane-targeted RFP mRNA.

### Heat-shock inducible expression of *cdkn1c*

*cdkn1c* was amplified from cDNA using the following primers: P57 forward: 5’-ATGGCAAACGTGGACGTATCAAGC-3’

P57 reverse: 5’-GCATGAAATTGCAAACCAAACTT-3’.

PCR product was cloned into the pCRII vector (TOPO-TA kit; Invitrogen), generating pCRII-cdkn1c. pCRII-cdkn1c was digested with EcoRI and then ligated into EcoRI-cut, shrimp alkaline phosphatase treated pSGH2 (Bajoghli et al., 2004), generating pKC040 to enable the expression of both the green fluorescent protein (GFP) and *cdkn1c* from bidirectional heat-shock elements. 15 ng/µl of either pSGH2 or pKC040 plasmid were injected into single-cell staged embryos and incubated at 25°C. To induce expression of GFP or GFP and *cdkn1c*, 28 hpf embryos were incubated at 38°C for 30 minutes, transferred to E3 with PTU, incubated until 3.5 dpf, and then fixed with 4% PFA for immunohistochemistry.

### Immunohistochemistry, imaging, and analysis

After fixation, wholemount embryos were either subjected to immunohistochemistry as previously described (Cerveny et al., 2010) or were cryoprotected in 15% and then 30% sucrose before being embedded in Optimal Cutting Temperature (OCT) resin and cut into 30 µm thick sections that were collected on charged glass slides (Polysciences, 24216) and stained with the following antibodies: beta-catenin (mouse, 1:250 dilution; Sigma, C7207); GFP (chicken, 1:250 dilution, Abcam, ab139709); RFP (rabbit, 1:500 dilution, MBL, PM005); PH3 (rabbit, 1:300 dilution, Millipore, 06-570); ZO-1 (mouse 1:100, Invitrogen, 339100). Nuclei were counterstained with DAPI (1 µg/ml from a 1 mg/ml stock in DMSO; Sigma) or sytox orange (1:10,000 dilution, Invitrogen).

All images were captured on a Nikon A1+ confocal with a long working distance 25X, 1.1 NA water immersion lens. To quantify clone size, stacks of confocal images were converted to Imaris (Bitplane) files and distinct clones were first manually contoured to generate distinct *cdkn1c* and GFP-positive surfaces. Surfaces were masked, generating a channel containing DAPI nuclei in each surface. Images were then batch processed to automatically count nuclei per clone. Clone sizes were exported to a spreadsheet, then graphed and statistically analyzed using JMP Pro 14 (Scintilla).

---

**Supplemental Figure 1.**
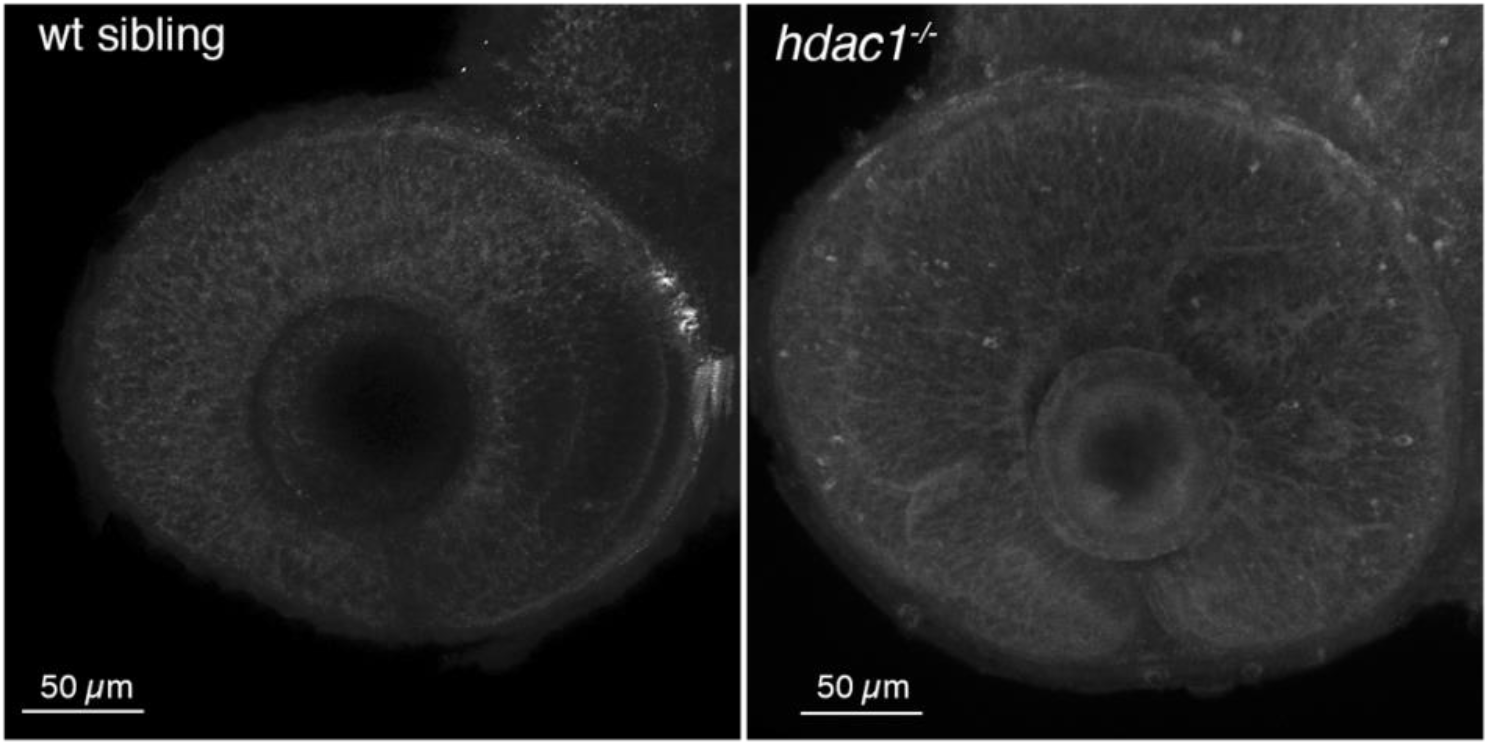
Retinal progenitor cells undergo apoptosis in *hdac1* mutant retinae. Wild-type siblings (left) and *hdac1* mutants (right) were fixed at ~3.5 dpf and then probed with activated caspase 3 antibody. Both representative images are maximum intensity projections. Note puncta scattered throughout hdac1 mutant but not in wild-type.

**Supplemental Figure 2.**
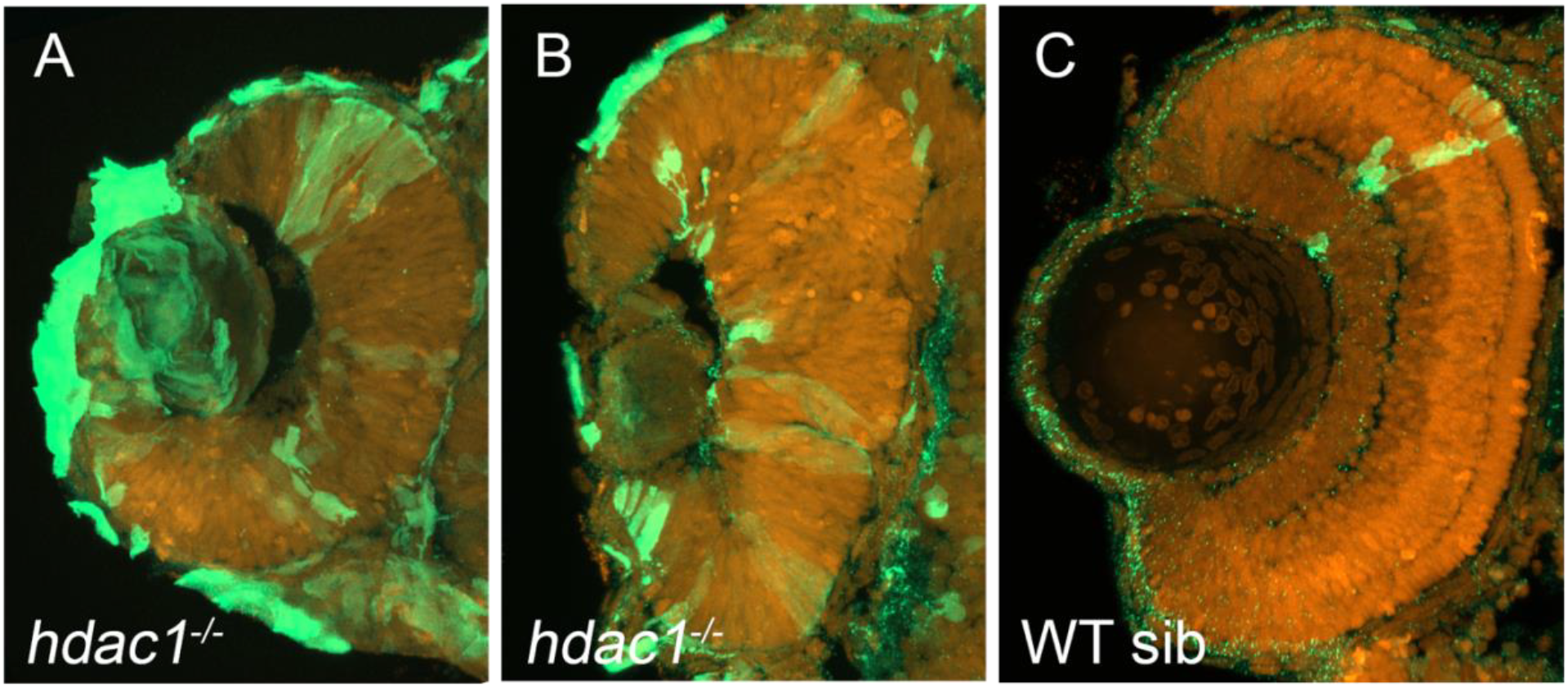
Inducing high levels of *cdkn1c* rarely trigger neuronal differentiation in *hdac1* retinal progenitor cells. Frontal cross-sections of retinae from same experiment shown in Figure 3; *cdkn1c*-positive clones were detected by immunohistochemistry (green) and nuclei stained with DAPI (red). *hdac1* mutant cells in *cdkn1c*-expressing clones (green) generally exhibit neuroepithelial morphology (A, B) consistent with proliferation and rarely show neuronal phenotypes (B, arrow heads) whereas age-matched wild-type sibling cells within *cdkn1c*-expressing clones (green) always exhibit neuronal morphology and lamination (C).

## Declarations

### Ethics approval

Adult zebrafish were bred and cared for with protocols approved by the Reed College IACUC.

### Consent for publication

All authors have read, provided feedback, and approved the manuscript.

### Availability of data and material

All data of this study are presented in this manuscript and any fish lines or other materials are available from the corresponding author or, with regard to mutant fish lines, from the Zebrafish International Resource Center (ZIRC; zebrafish.org).

### Author contributions

This study was conceived as an outgrowth of a project that KLC began as a post-doc in the lab of Stephen W. Wilson at University College London with consultation from MV. KLC, AVD, IT, HB, OH, and DBL performed experiments and made figures. MV supplied the *gins2* morpholino and additional data before publication. KLC wrote the manuscript with input from all authors.

## Acknowledgements

We thank all members of the Cerveny lab and especially Steve Wilson where the idea of this project took root. We also thank the ZIRC for serving as a repository for all mutant lines and advice about fish rearing.

